# Characterization of the different behaviours exhibited by juvenile grey mullet (*Mugil cephalus*) under rearing conditions

**DOI:** 10.1101/2021.02.23.432396

**Authors:** Jessica A. Jimenez-Rivera, Anaïs Boglino, Joel F. Linares-Cordova, Neil J. Duncan, María de Lourdes Ruiz-Gómez, Sonia Rey Planellas, Zohar Ibarra-Zatarain

**Affiliations:** Posgrado en Ciencias Biológico Agropecuarias, Universidad Autónoma de Nayarit, 63155 Tepic, México; CONACYT-UAN-Nayarit Centre for Innovation and Technological Transference (CENITT), Av. E González s/n. CP 63173. Tepic, México; Nayarit Centre for Innovation and Technological Transference (CENITT), Av. E González s/n. CP 63173. Tepic, México; Posgrado de Ciencias Agropecuarias, Colegio de Ciencias Agropecuarias, Universidad Autónoma de Sinaloa, Km 17,5, CP 8000 Culiacán, México; IRTA, Sant Carles de la Ràpita, Carretera de Poble Nou, km 5.5, 43540 Sant Carles de la Ràpita, Tarragona, Spain; Laboratorio de Ecología y Conducta, Facultad de Ciencias, Universidad Autónoma del Estado de México, Toluca, Estado de México, México; Institute of Aquaculture, Faculty of Natural Sciences, University of Stirling, FK9 4LA, Stirling, Scotland, UK

**Keywords:** welfare, behaviour, grey mullet, rearing conditions, ethogram

## Abstract

This research described the common behaviour of grey mullet (*Mugil cephalus*) under rearing conditions. The different behaviours exhibited by mullets were videorecorded with submersible cameras installed inside of three tanks. A total of 690 minutes per day (07:30 - 18:30 hours) were recorded per tank during a week. Afterwards, an ethogram was elaborated to organize the different behaviours exhibited by juvenile *M. cephalus*, with two general categories: a) locomotion, including three different observed behaviours (resting, swimming and fast swimming) b) feeding, including three behaviours (surface feeding, bottom feeding and rubbing). The video recordings showed that *M. cephalus* is a species with a constant locomotion associated to feeding, since they showed constantly movement during most of day light period the opposite to dark periods. Mullets were observed to be a non-aggressive fish species, due to the absence of dominance and aggression towards conspecifics, resulting in a high predisposition for adaptation to captivity. Finally, behavioural frequencies of grey mullet’s juveniles were not significantly different among the three tanks for most of the behavioural variables analysed (*p*>0.05) except for the variable bottom feeding (*p*=0.02). Results from this study could be of interest to the aquaculture industry to implement protocols and to optimize rearing techniques for the production of grey mullet.

## Introduction

*Mugil cephalus*, commonly known as grey mullet, is a cosmopolitan fish species. It is found in all oceans and in a large variety of aquatic environments. It has a high tolerance to different environmental conditions that includes a wide range of temperatures and salinities (Saleh, 2008). These eurytopic attributes of grey mullet, in combination with the foraging feeding habits and fast-growing rate (~0.70 kg per year), enable this species to be considered for both freshwater and marine aquaculture (Whitfield et al., 2012; FAO, 2020a). Overall, grey mullet has been used as a model in different research areas, such as ecotoxicology, population dynamics, parasitism and gametes cryo-preservation (Chao & liao, 2001; Mahanty *et al*., 2011; Crosetti & Blaber, 2015; Colin *et al*., 2020). However, the interest for human consumption has increased during the last decade and data provided by FAO (2020b) showed that grey mullet fisheries has increased in the last decade from 101,182 tons in 2008 to 130,233 tons worldwide in 2018. This increasing number of captures has led to a growing interest by the aquaculture industry for developing techniques and protocols to produce fingerlings in captivity. In this context, mullet aquaculture production was of 13,681 tons in 2016, where Egypt being the main producer, followed by the Republic of Korea, Italy, the Chinese Province of Taiwan and Israel (Crosetti, 2015; FAO, 2020a). Likewise, in some countries the polyculture practices of *Mugil cephalus* (Soto, 2009) with other species such as *Penaeus vannamei* (Hosseini Aghuzbeni *et al*., 2017), *Penaeus monodon* (Mondal *et al*., 2020) turn out to be a sustainable alternative.

The economic importance of *Mugil* is based on its consumption and its cost varies from one region to another, relying on several factors; for example, female’s grey mullet gonads, also known as caviar or bottarga, could reach 65 € per kg in the European markets (Aldana, 2015; Rodríguez, 2018). Therefore, significant advances have been made on nutrition, growth, larval culture and reproduction of individuals reared in captivity (Martínez *et al*., 2019; Talukdar *et al*., 2020; Besbes *et al*., 2020; Ramos-Júdez *et al*., 2021). Furthermore, several studies have demonstrated that grey mullet can also be an adequate candidate species for mariculture which can contribute to food production and the reduction of fishing impact (Saleh, 2008; Robles & Mylonas, 2017).

For domesticating and rearing a fish species, welfare is important, since confinement conditions trigger physiological and behavioural responses which impact growth performance, diseases outbreaks, reproduction, among other factors (Ashley, 2007). It has also been recognized that most of the typical behaviours exhibited by organisms in captivity, such as: feeding, swimming, sociability, dominance, reproduction, etc., are related to environmental stimuli and aquaculture management practices (Rowland, 1999; Lall & Tibbetts, 2009; Baran & Streelman, 2020). Additionally, the behavioural responses of individuals under rearing conditions are often used as operational welfare indicators, since they might indicate potential stressful situations of individuals in their environment (Huntingford *et al*., 2006). Hence, understanding the behaviour of cultured species can be an early warning of alterations in animal stress status and health, and a useful tool to provide adequate environmental conditions and facilities promoting welfare of the reared organisms (Kristiansen *et al*., 2004; Saraiva *et al*., 2019).

In behavioural studies, ethograms provide reliable information about the behavioural responses of animals in their environment. According to McDonnell & Poulin (2002), an ethogram could be defined as a formal description of a species behavioural repertoire or a major segment of it. It may be a complete list of all behaviours or it may focus on particular functional classes of behaviours. Therefore, an ethogram shows the actions, interactions and overall activity typically performed by animals in the wild and such responses are expected to be replicated in captivity (Marsh & Hanlon, 2004). Currently, ethological studies of fish in captivity includes the analysis of behavioural functional categories, such as reproductive behaviours (Ibarra-Zatarain & Duncan, 2015) or feeding characteristics (Huntingford, 2004; Carvalho *et al*., 2007). However, studies that analyse in detail the common behaviour exhibited by the organisms under captivity conditions are scarce or inexistent for many species that are currently produced in aquaculture (Lahitte *et al*., 2002; Bolgan *et al*., 2016). In the case of *M. cephalus*, there are no studies describing the common behaviour of this species in the wild or under rearing conditions. Therefore, it is relevant to describe the behaviour of this species in captivity, since it represents a forecasting tool to evaluate the preferences and requirements of the animals, to provide adequate management protocols and facilities that improve their welfare (Castanheira *et al*., 2017; Saraiva *et al*., 2019). Hence, the aim of this study was to generate an ethogram that described the common behavioural exhibited by juveniles of grey mullet *Mugil cephalus* in rearing conditions.

## Materials and methods

### Ethic statement

The number, handling and manipulation of the organisms used in this study were established following the criteria of the National Centre for the Replacement, Refinement and Reduction in Animals in Research (NC3Rs, U.K.). Locally, the protocol for handling and use of animals was authorized by the Bioethics Commission of the State of Nayarit, Mexico (permit number CEBN / 05/2017).

### Collection and maintenance of organisms

Fish were captured from the wild on the pacific coast, Mazatlán, Mexico, in September 2019. A total of 300 fish were caught (average weight and length 30.2 ± 6.9 g and 15.1 ± 1.2 cm, respectively). Fish were transported to the Nayarit Centre for Innovation and Technological Transference (CENITT-UAN) in Tepic, Nayarit. Individuals were acclimated in two 500L rectangular tanks (100 x 140 x 55 cm) connected to a recirculation system (RAS). Once acclimation was completed (45 days), a total of 36 fish were randomly selected and transferred to three 220 L rectangular tanks (80 x 68 x 48 cm) to reach a final density of 12 fish per tank during the experiment. Water parameters were maintained as follows: temperature: 25-27°C, salinity: 27-29 ppm, pH: 6-7 and oxygen 5-7 mg / L. All water parameters were monitored daily in the morning. Photoperiod was adjusted to follow the natural seasonal cycle (Light: Dark, 11: 13) by using an automated external dimmer (MyTouchSmart, General Electric^®^) turning on-off white lamps (OSRAM 85Watts) at 08:00 – 19:00 hours during the experiment. Mullets were fed to satiety with a commercial diet designed for marine fish (specifications: floating pellets, 55% crude protein, 3.00 mm; Skretting®, The Netherlands). Tanks were siphoned each day, 30 min before the first feeding and 30 min after the last feeding, to remove the remains of food and faeces and to maintain adequate water quality conditions.

### Description of behaviour in grey mullet (*Mugil cephalus*)

#### Data collection

A high-definition camera system (Swann/2K Series-1080p) was installed in three tanks. Each camera was positioned on the lateral wall of the tank, 10 cm below the water surface in order to capture more than 90% of the total area of the tank. Video recording started at 07:30 and finalized at 19:00 h every day for one week. The selection and quantification of behavioural variables were based on focal observations of the video recordings performed by three distinct observers, whom reported traits of the behavioural repertoire of the organisms. It included feeding habits, swimming and overall activity at the beginning and end of seven consecutive days, following the recommendations of Bolgan *et al*. (2016). A total of 690 minutes per day were recorded from each tank, comprising 90 video analysis per experimental unit (Mas-Muñoz *et al*., 2011; Ibarra-Zatarain & Duncan, 2015; Thomsen *et al*., 2020).

#### Behavioural variables

The behavioural variables analysed and selected were based on Altman (1974), and were grouped in two categories: 1) *locomotion* and 2) *feeding*. For the first category (Table 1), three behaviours were evaluated and quantified: i) resting, ii) swimming and iii) fast swimming. For the second category (Table 1), three behavioural variables were evaluated: i) surface feeding, ii) bottom and feeding iii) rubbing. These behavioural variables were selected and adapted from previous descriptive studies on behaviour with other marine and freshwater fish species, such as the bicolour damselfish *Eupomacentrus partitus* (Myrberg, 1972), common sole *Solea solea* (Mas-Muñoz *et al*., 2011), some reef fishes (Pink & Fulton, 2014), gilthead seabream *Sparus aurata* (Ibarra-Zatarain & Duncan, 2015) and artic charr *Salvelinus alpinus* (Bolgan *et al*., 2015).

**Table 1.** Description of the behavioural patterns observed in *Mugil cephalus*.

| Behaviour pattern | Description |
| --- | --- |
| **Resting** | Maintained a stationary position or slow swimming |
| **Swimming** | Fluid and constant swimming in group or individual |
| **Fast swimming** | Subcarangiform locomotion |
| **Surface feeding** | Capture pellets on the tank surface with multidirectional movements that allow passage to other fish for that can feed |
| **Bottom feeding** | Complete exploration of the tank bottom until the intake of all the food offered |
| **Rubbing** | The fish rub their mouths against the tank surfaces in a vertical position with shaky movements type zigzag |

#### Statistical analysis

Statistical analyses were performed using IBM SPSS Statistics 25 software. Data were checked for normality and homoscedasticity with a Kolmogorov-Smirnov test and a Levene’s test, respectively. A multivariate analysis of variance (MANOVA) was performed to assess the uniformity in the behaviours analysed among the three tanks where the cameras were installed. Additionally, a Tukey’s post-hoc test was performed on data when significant differences were detected among tanks. A 95% confidence interval (*p*=0.05) was set for all analyses.

## RESULTS

### General behavioural observations

*Mugil cephalus* exhibited a stationary swimming behaviour or resting during the periods with no illumination before and lights were switched on and off (07:30 to 08:00 and 18:30 to 19:00 hours, respectively). Locomotor activity of grey mullet juveniles increased gradually during the day and this behaviour was similar in all tanks throughout the experiment. Additionally, mullets were observed to be a social, non-aggressive and highly active fish species.

### Description of behaviour

#### Locomotion

##### Resting

This behaviour was mainly characterized by stationary swimming, where fish pectoral and tail fins remained close to their body with slow undulations. Fish maintained a stationary position in the water column; however, in a few occasions, they performed slow movements. Moreover, fish did not exhibit social interactions, since they occupied different positions in the water column (Fig. 1-A, Table 1). This inactivity occurred always in the absence of light, before 08:30 and after 18:40 hours and could be described as fish sleep (Keene & Appelbaum, 2019).

**Figure 1.**
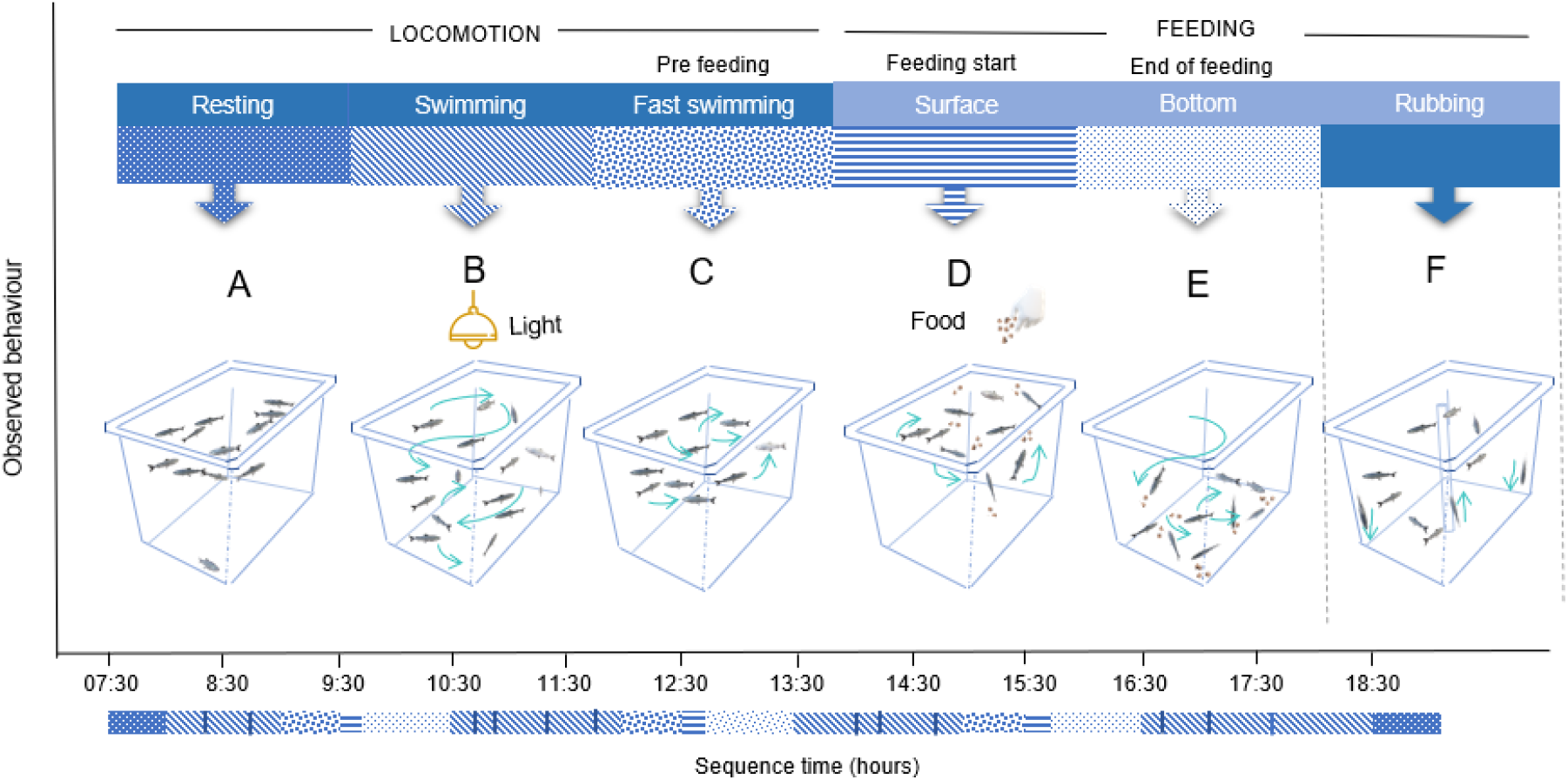
Behavioural patterns observed in *Mugil cephalus* in captivity. The color variation represents each behavioural pattern exhibited by the fish in chronological order during the observation period (7:30 - 18:30 hours).

##### Swimming

This behaviour was characterized by active swimming, in which fish swam at a constant speed throughout the tank. Additionally, mullets formed small groups or fish shoals with some individuals swimming independently. While fish were swimming, they took different directions and interacted with other individuals in repeated occasions. Moreover, no aggressive behaviours such as chases, bites, fin erections, etc. were detected. This behaviour was observed in daylight hours, between 08:45 and 18:30 hours (Fig. 1-B, Table 1).

##### Fast swimming

This is defined as high fish activity or fast swimming, in which fish presented a subcarangiform locomotion consisting in constant undulations of the posterior part of the body, accompanied by movements of the tail fin and with or without extensions of pectoral fins. Fish showed prolonged movements from one point to another in the water column, with constant interactions between fish, but with no aggression. This behaviour occurred approximately 30 minutes before a feeding event (Fig. 1-C, Table 1).

#### Feeding

##### Surface feeding

When feed was offered, fish quickly approached and ingested a maximum number of pellets. However, it was also noted that after capturing the pellets, fish frequently spitted it into the water. During the feeding frenzy, mullets showed multidirectional movements around the food. In addition, a cooperative event between fish was observed in which animals that first consumed pellets, moved immediately to another point in the water column, allowing other individuals to rise at the surface to feed. Besides, no signs of aggression or dominance over food were detected (Fig. 1-D, Table 1).

##### Bottom feeding

After the food was sank, grey mullet began a second feeding event in which they consumed the food in the bottom of the tank in a vertical position. In this context, fish were observed to explore the bottom of the tank searching for uneaten pellets, and constantly interact with other fish, but no aggressive behaviour were neither observed (Fig. 1-E, Table 1). Unlike surface feeding, food consumed at the bottom of the tank was swallowed and not spited into the water.

##### Rubbing

This behavioural was distinctive of fish by rubbing their mouths against the walls or bottom of the tank doing zigzag movements. While performing this behaviour, fish maintained a vertical position with the pectoral fins close to the body. Rubbing was observed individually or in group (less than 60% of the fish) in different occasions along the day. (Fig. 1-F, Table 1).

#### Comparison of behavioural frequencies

No statistical differences (*p*>0.05) were detected in the frequencies of the six behavioural variables analysed among the tanks (Table 2). Specifically, no significant differences were detected for resting (*p*=0.124), swimming (*p*=0.064), fast swimming (*p*=0.519), surface feeding (*p*=0.122) and rubbing behaviour (*p*=0.053). Fish from tank 2 presented significant higher frequencies in bottom feeding than fish from tank 3 (*p*=0.026). Therefore, the results suggest that grey mullet were successfully adapted to captivity, since their behavioural tendencies were similar among tanks.

**Table 2.**
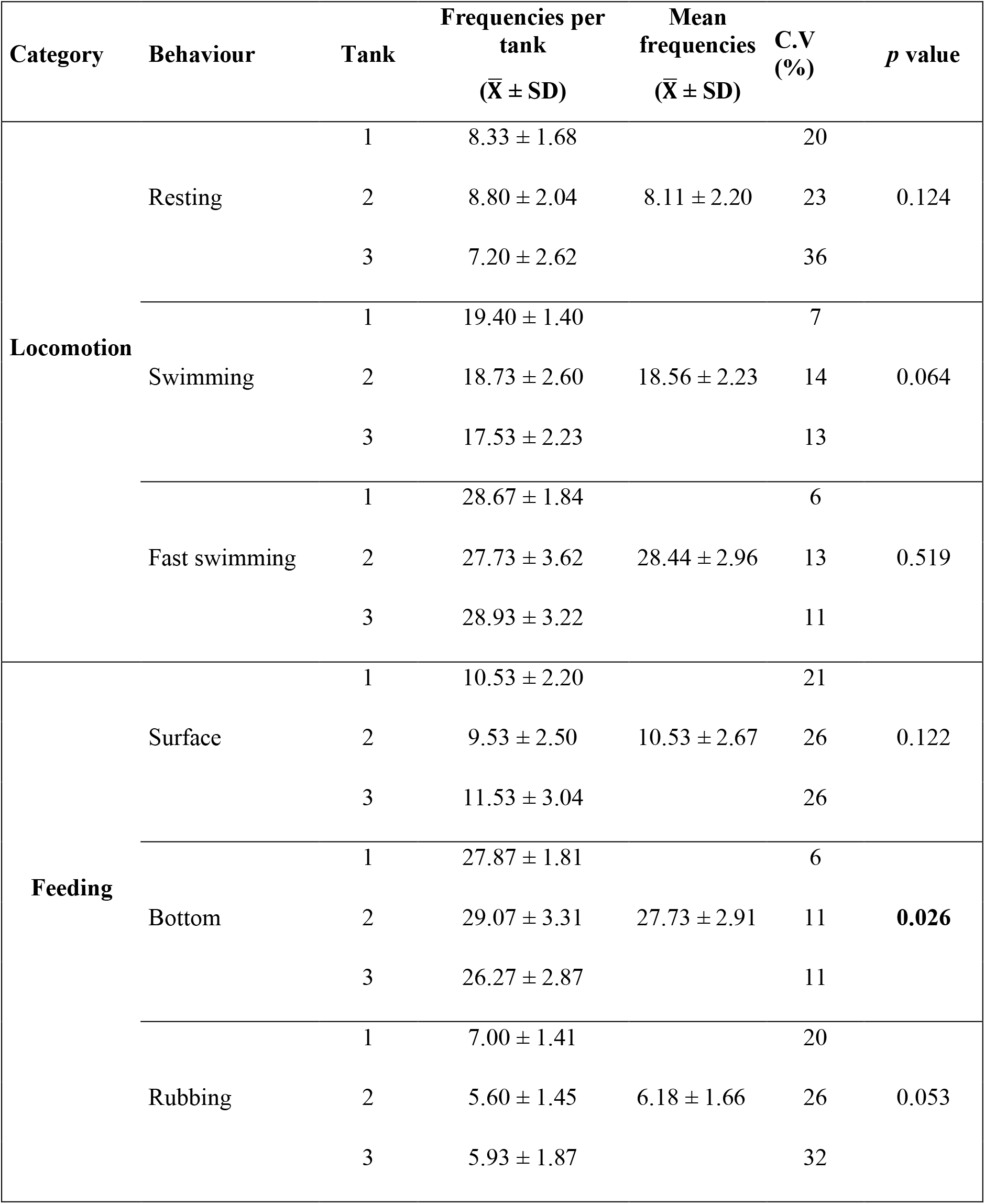
Frequencies observed for the different behaviour patterns expressed in juvenile *Mugil cephalus* in captivity. 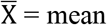 observed frequencies ± standard deviation; CV = coefficient of variation. Values in bold represent significant differences from the MANOVA test.

| Category | Behaviour | Tank | Frequencies per tank<br>($\bar{X} \pm SD$) | Mean frequencies<br>($\bar{X} \pm SD$) | C.V<br>(%) | *p* value |
| --- | --- | --- | --- | --- | --- | --- |
| Locomotion | Resting | 1 | 8.33 ± 1.68 | | 20 | |
| | | 2 | 8.80 ± 2.04 | 8.11 ± 2.20 | 23 | 0.124 |
| | | 3 | 7.20 ± 2.62 | | 36 | |
| | Swimming | 1 | 19.40 ± 1.40 | | 7 | |
| | | 2 | 18.73 ± 2.60 | 18.56 ± 2.23 | 14 | 0.064 |
| | | 3 | 17.53 ± 2.23 | | 13 | |
| | Fast swimming | 1 | 28.67 ± 1.84 | | 6 | |
| | | 2 | 27.73 ± 3.62 | 28.44 ± 2.96 | 13 | 0.519 |
| | | 3 | 28.93 ± 3.22 | | 11 | |
| Feeding | Surface | 1 | 10.53 ± 2.20 | | 21 | |
| | | 2 | 9.53 ± 2.50 | 10.53 ± 2.67 | 26 | 0.122 |
| | | 3 | 11.53 ± 3.04 | | 26 | |
| | Bottom | 1 | 27.87 ± 1.81 | | 6 | |
| | | 2 | 29.07 ± 3.31 | 27.73 ± 2.91 | 11 | **0.026** |
| | | 3 | 26.27 ± 2.87 | | 11 | |
| | Rubbing | 1 | 7.00 ± 1.41 | | 20 | |
| | | 2 | 5.60 ± 1.45 | 6.18 ± 1.66 | 26 | 0.053 |
| | | 3 | 5.93 ± 1.87 | | 32 | |

## Discussion

This study described for the first time the common behaviour of juvenile *M. cephalus* under captive. Overall, mullets are individuals with most activity during daytime due their high rates of locomotion, swimming, social interactions during the day, which decrease immediately after lights were turned off, as a transition to resting behaviour. In this context, Helfman (1986) suggested that the behaviour in animals, such as: feeding, breeding, aggregations and resting, are influenced by the artificial light alternation (light and dark) as was the case for this fish species.

### Locomotion

The locomotor activity of grey mullet started with a resting behaviour or slow swimming, registered during the first minutes of the day (between 07:30 to 08:00 hours) and when lights were turned off (between 18:30 to 19:00 hours). Bolgan *et al*. (2016) described a similar resting behaviour in the arctic charr (*Salvelinus alpinus*) held in captivity. Moreover, authors classified resting behaviour as a state of inactivity, in which fish held a stationary position most of the time with no forward locomotion. Similarly, Ibarra-Zatarain & Duncan (2015) reported that gilthead seabream tended to swim slowly either alone or in small groups around the tank during early morning (08:30 hours). Likewise, Park *et al*. (2018) reported the same resting behaviour in the Korean endemic cobitid (*Iksookimia hugowolfeldi*) in the wild. Killen *et al*. (2016) have suggested that resting behaviour is associated to energy conservation. Also, Zimerman *et al*. (2008) and Elbaz *et al*. (2013) have described that prolonged periods of behavioural quiescence could be defined as sleep in fish. Those authors pointed out that when fish decrease locomotor activity and metabolic rate in order to save energy. Therefore, energy is conserved to be allocated for functional activities, such as feeding, growing or reproduction, which are important biological parameters for aquaculture.

The second locomotor pattern analysed, swimming behaviour, was the most common in grey mullet juveniles and it was characterized by a constant movement of the fish day light hours. According to different studies, this conduct has been typically described for other fish species and is linked to their physiology. For example, Farwell & McLaughlin (2009) and Brownscombe *et al*. (2017) suggested that constant swimming behaviour is adaptive to environment and might be linked to biological functions such as: metabolism, respiration and digestion. A similar conduct to that described in the present study was reported by Ibarra-Zatarain & Duncan (2015), whom observed that gilthead seabream (*Sparus aurata*) had a swimming activity characterized by a constant swimming speed in all fish. Furthermore, swimming behaviour in white mullet (*Mugil curema*) in their natural environment (Carvalho *et al*., 2007) coincided with the conduct reported in the present study, is also characterized by constant swimming through the day. Downie *et al*. (2020) mentioned that constant swimming behaviour denotes optimal welfare and a good physiological condition.

Regarding fast swimming behaviour, it has been documented that locomotor activity tends to increase when it is associated to behaviours such as: feeding, foraging and mating in captivity or predator avoidance and migration in the wild (Brownscombe *et al*., 2017). Therefore, the fast swimming behaviour exhibited by *M. cephalus* in captivity, may be related to an anticipatory activity similar to what happens in other aquaculture species. For example, gilthead seabream (Montoya *et al*., 2010) and tambaqui (*Colossoma macropomum*) (Fortes-Silva *et al*., 2018) showed an increased or anticipatory swimming activity before feeding. In a different context, other studies have reported that fast swimming, could be associated fast growing fish (Palstra *et al*., 2010), as they present optimal muscular-skeletal development and osmoregulation (Huntingford and Kadri, 2013) and exhibits stress resistance (Martins *et al*., 2012).

### Feeding

Mullets showed two distinctive feeding behaviours. First, individuals showed a preference to swallow pellets in the surface of water. This initial feeding reaction could represent an intuitive strategy securing food (Montoya *et al*., 2010); as reported in European seabass (*Dicentrarchus labrax*) (Azzaydi *et al*., 1998) and *Parachaeturichthys ocellatus* (Paniker, 2020). Moreover, it is known that feeding fish regularly, like in aquaculture rearing conditions, induces an internal mechanism of synchronization with food. Additionally, Lall & Tibbetts (2009) proposed that the feeding behaviour in fish is associated to cognition, similarly to birds and mammal. Therefore, it is possible that grey mullet (*Mugil cephalus*) adapts its behaviour to fixed schedules of feeding in captivity conditions.

After grey mullets consumed the pellets in the water surface, it was observed that some individuals frequently spitted out the pellets that they have just swallowed resulting in approximately 60% of the offered food sinking to the bottom of the tank when second feeding started. This behaviour lasted until fish consumed all the pellets and it was accompanied by exploration and social interactions. Islam *et al*. (2009) suggested that grey mullet is a bottom feeder, that exhibits herbivore preferences, based on gut content, which may explain the presence of bottom feeding in captivity. Similar results have been found by Anders *et al*. (2017) and Park *et al*. (2018) in cod (*Gadus morhua*) and the Korean endemic cobitid, suggesting that this preference to feed is associated with their natural feeding habits. When analysing the frequencies of this behavioural variable, significant differences were detected among tanks. This variation could be related to feeding practices, since fish were fed *ab-libitum* and, thus, it is possible that fish did not receive the same amount of feed. To confirm this previous assumption, it will be important to evaluate different food rations for this fish species, as suggested by Wassef *et al*. (2001) who reported that 4% of total biomass in food, might result in similar behavioural frequencies.

On the other hand, rubbing behaviour has been related to the lifestyle and feeding strategy of this fish species (Almada *et al*., 1999), and similar observation was reported in the korean endemic cobitid. This fish species showed a zig zag swimming behaviour which was associated with their habitat and feeding behaviour (Park *et al*., 2018). Additionally, Islam *et al*. (2009) described that mullet feed in the wild mainly with algae and detritus; thus, the adaptation of feeding behaviour in the Mugilidae family is instinctively based on searching for food at the bottom or edges by rubbing its body to graze and ingest phytoplankton and other detritus (Bowen, 1984). Therefore, by showing this kind of behaviour, grey mullet may be adapting to captivity, hence, this behavioural response can be used as a measurement of optimal performance of the species.

Lastly, mullet seemed to be a social species, due to the absence of aggression or the establishment of social hierarchies. Additionally, grey mullet juveniles exhibited a cooperative behaviour when consuming food swimming in a synchronized way, allowing all individuals to eat. Wey *et al*. (2008) described that social animals live and interact together, forming complex social relationships and structures that can have advantageous to obtain resources (i.e. food, feeding areas, others). Unlike, in other reared species such as the rainbow trout there is a marked competition for resources, (Øverli *et al*., 2004) or the presence of dominance in sole, conferring an advantage of some fish over others access to the food (Fatsini *et al*., 2017). Therefore, the absence of hostile behaviours in the grey mullet could be considered as an important factor for the aquaculture of this fish species.

## Conclusion

In conclusion, behaviours are responses exhibited by fish in their environment to access to resources, which allow them to meet their basic requirements and survive (Føre *et al*., 2018). Considering that welfare of reared animals generally leads to a production of quality, it is relevant to know the expression of natural behaviours as a forecasting model of welfare status and physiological performance (Huntingford, 2004). Thus, this research described the typical behavioural repertoire of mullet juveniles (*Mugil cephalus*), where all the functional categories observed in rearing conditions were described, that is, the usual behaviour observed in chronological order in a production farm. Moreover, traits such as cognition and social behaviour may pose the attraction for mullet as an attractive aquaculture species in Mexico. Thus, this study contributes to the knowledge of the species for a correct management of the biological processes and to implement rearing protocols that will improve aquaculture production of a species of economic interest. Moreover, this study represents a starting point for any behavioural study in mullet and a reference for comparison with others studies in the wild and under stress conditions for this species.

## Notes

### Competing Interest Statement

The authors have declared no competing interest.

